# Fractal Genomics of *SOD1* Evolution

**DOI:** 10.1101/2020.05.14.097022

**Authors:** Mohammad Saeed

## Abstract

To understand the fundamental processes of gene evolution such as the impact of point mutations and segmental duplications on statistical topography, Superoxide Dismutase-1 (*SOD1*) orthologous sequences (n=50) were studied. These demonstrated scale invariant self-similarity patterns and long-range correlations (LRC) indicating fractal organization. Phylogenetic hierarchies changed when *SOD1* orthologs were grouped according to fractal measures, indicating statistical topographies can be used to study gene evolution. Sliding window k-mer analysis showed that majority of k-mers across all *SOD1* orthologs were unique, with very few duplications. Orthologs from simpler species contributed minimally (<1% of k-mers) to more complex species. Both simple and complex random processes failed to produce significant matching k-mer sequences for *SOD1* orthologs. Point mutations causing amyotrophic lateral sclerosis did not impact the fractal organization of human *SOD1*. Hence, *SOD1* did not evolve by a patchwork of repetitive sequences modified by point mutations. Instead, this study proposes that *SOD1* gene sequences evolved by regulated interweaving of unique oligomer sequences that led to LRC, signifying convergent evolution.

**Summary Statement:** *SOD1* has long-range correlations which resulted not from point mutations, segmental duplications or patching together sequences from simpler organisms. Instead, *SOD1* underwent convergent evolution by repeated unique sequence assemblies.

## Introduction

The extraordinary complexity of biological systems is generated by repeating patterns coded by recursive processes, which can be quantified by fractal analysis. These irregular processes that appear to be random are in fact governed by rules of chaos – self-similarity, sensitivity to initial conditions and long-range order over multiple scales of space and time (Mandelbrot, 1982; Peitgen et al. 2004). Understanding the basic paradigm of biological rhythms may help predict their occurrence and eliminate abnormality. The randomness incorporated in genomes is mathematically chaotic with multiple statistical topographies such as periodicities and long-range correlations (LRC), resulting in a fractal organization (Peng et al., 1992; Voss, 1992; Buldyrev et al., 1993; Albrecht-Buehler, 2012). Studying genomic organization will not only provide insight into evolutionary biology but also disease mechanisms, potentially yielding predictive tools for clinical medicine.

Random mutation and natural selection are known to be two forces that have shaped evolution. The degree to which simple random processes influence the final development of natural genomic sequences is of critical importance to biology and medicine. In this paper we attempt to understand how functional genomic sequences developed from short strands of randomly assorted nucleotides. The Superoxide Dismutase-1 (*SOD1*) gene is a fundamental component of redox systems across a broad range of species. Moreover, *SOD1* mutations cause amyotrophic lateral sclerosis (ALS) (Saeed et al., 2009). Hence, *SOD1* was chosen to study the evolutionary processes and the impact of sequence changes (point mutations and segmental duplications) on genomic organization and phylogeny.

Since the time LRC have been described in DNA, questions have arisen about their purpose and mechanism(s) of development (Peng et al., 1992; Maddox, 1992; Moreno et al., 2011). It was proposed that LRC occur due to segmental duplication and modified by point mutations (Li and Kaneko, 1992; Messer et al., 2005). Though these processes can theoretically produce LRC, this study shows that they play minor roles if any in the evolution of natural genomic sequences such as *SOD1*. The data presented here shows that short range correlations are present as well (in small sized sequences from lower organisms). Finally, this paper shows that *SOD1* did not evolve by patching together sequences from simpler organisms. Instead, *SOD1* underwent convergent evolution by repeated unique assemblies of oligomer sequences, consequently leading to the development of LRC.

## Results

### Sequence correlations exist

DFA α ranged from 0.45 in *Emiliania huxleyi* (Gene ID: 17260763) to 0.74 in *Homo sapiens* (Gene ID: 6647). As shown in Table S1, 46 *SOD1* orthologs showed LRC (α > 0.5). There was moderate degree of concordance between DFA and RDA (R^2^ = 0.66) indicating that the two algorithms measure fractality differently. Neither measure correlated with gene length (DFA R^2^ = 0.48; RDA R^2^ = 0.41) or taxonomic order (DFA R^2^ = 0.49; RDA R^2^ = 0.38). Small gene sequences from simpler organisms such as *Caenorhabditis elegans* (174141) and *Apis mellifera* (409398) with length of ~ 1200 nucleotides showed correlations (α = 0.60 and 0.67; D = 1.47 and 1.34 respectively). Hence, short range correlations were present as well (Table S1). Exons of Human *SOD1* (NM_000454.5) also showed some degree of LRC (alpha = 0.533, D=1.33).

### Diagrammatic Self-similarity of *SOD1* sequences

To be consistent with fractals, *SOD1* sequences should form scale invariant patterns within a given *SOD1* ortholog (Albrecht-Buehler, 2012). These patterns may not be exact, however, their features should repeat across a range of sizes. Hence, the *SOD1* sequences were graphed in proportional window sizes to visually evaluate self-similarity, as in zoomed-in photographs. To allow uniform comparison of patterns across different sizes the *SOD1* sequence of length 2^n^ nucleotides, was allowed to weave across the plane in all four directions except that it wrapped up when it approached the window borders. Moreover, the window sizes were determined by the RDA principle to be 2^n+3^ to allow sufficient space for uniform pattern development. This photographic method is intuitively sensitive as pattern discernment is a fundamental ability of human vision. As shown in Figure 1, there is sufficient self-similarity (3-5 orders of magnitude) in the *SOD1* sequences to be considered as fractals. Further examples of the various degrees of self-similarity are depicted in Figure S1 and a complete list of images of *SOD1* orthologs can be viewed at: GeneFractals SOD1 Images.

**Figure 1.**
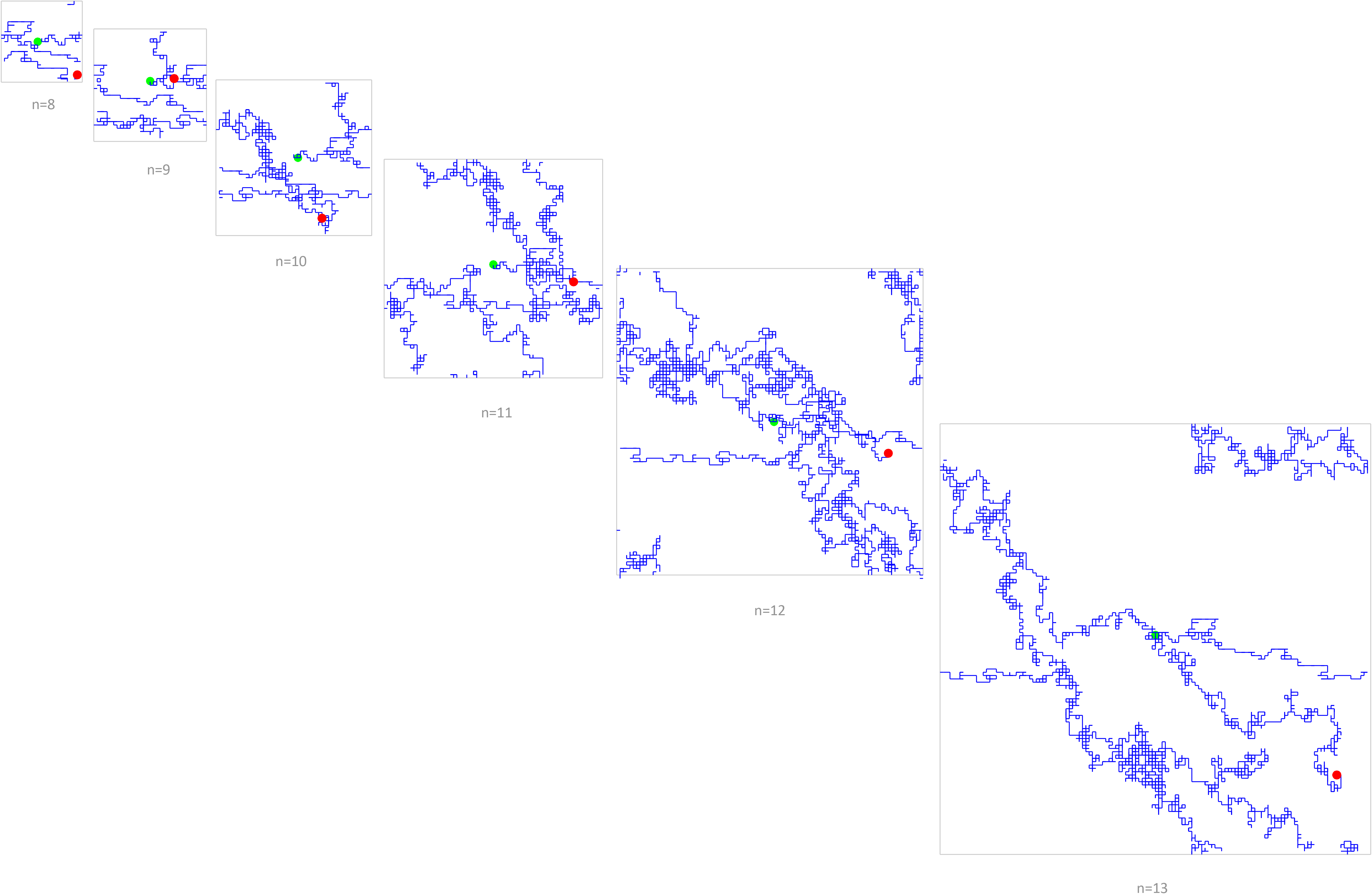
Fractal patterns of *SOD1* genetic sequences. *SOD1* sequence of *Gallus gallus* (Gene ID: 395938) of length up to 2^n^ nucleotides was graphically displayed, as described in the methods section. Self-similarity across multiple scales of length is visualized as in zoomed-in photographs. Fractal measures for *Gallus gallus SOD1* were α = 0.598 and *D* = 1.27.

### Phylogenetic sequence reconstruction of *SOD1*

Frequency matching of k-mer oligonucleotides showed that lower organisms contributed minimally (<1%) to genomic sequence of higher organisms (Figure 2). There was high frequency matching of 10-mers in specific groups of organisms. These groups somewhat resembled the DFA categories. The best example was that *Apis mellifera* (Gene ID: 409398) shared 7% of k-mers with *Phascolarctos cinereus* (Gene ID: 110214887) in comparison to the usual 1% matching (Table S2), with both having an α=0.67. Contribution to *Homo sapiens*’ *SOD1* (6647) by its orthologs was <10% by all organisms except two (*Macaca mulatta*: 574096 and *Pan troglodytes*: 449637). Half the organisms investigated (n=24) contributed <1% of 10-mers to human *SOD1*. Of the 226,923 total oligomers in 50 orthologs, 96.3% k-mers occurred only once in an organism. When all oligomers were pooled, 69% of these occurred only once across all phylogeny. On average <5% 10-mers (overall mean 2.1 ± 1.34%, range 0.32 – 4.79%) matched across all orthologs, indicating that *SOD1* gene in higher organisms was not derived by patching together sequences from taxonomically lower organisms (Table S2).

**Figure 2.**
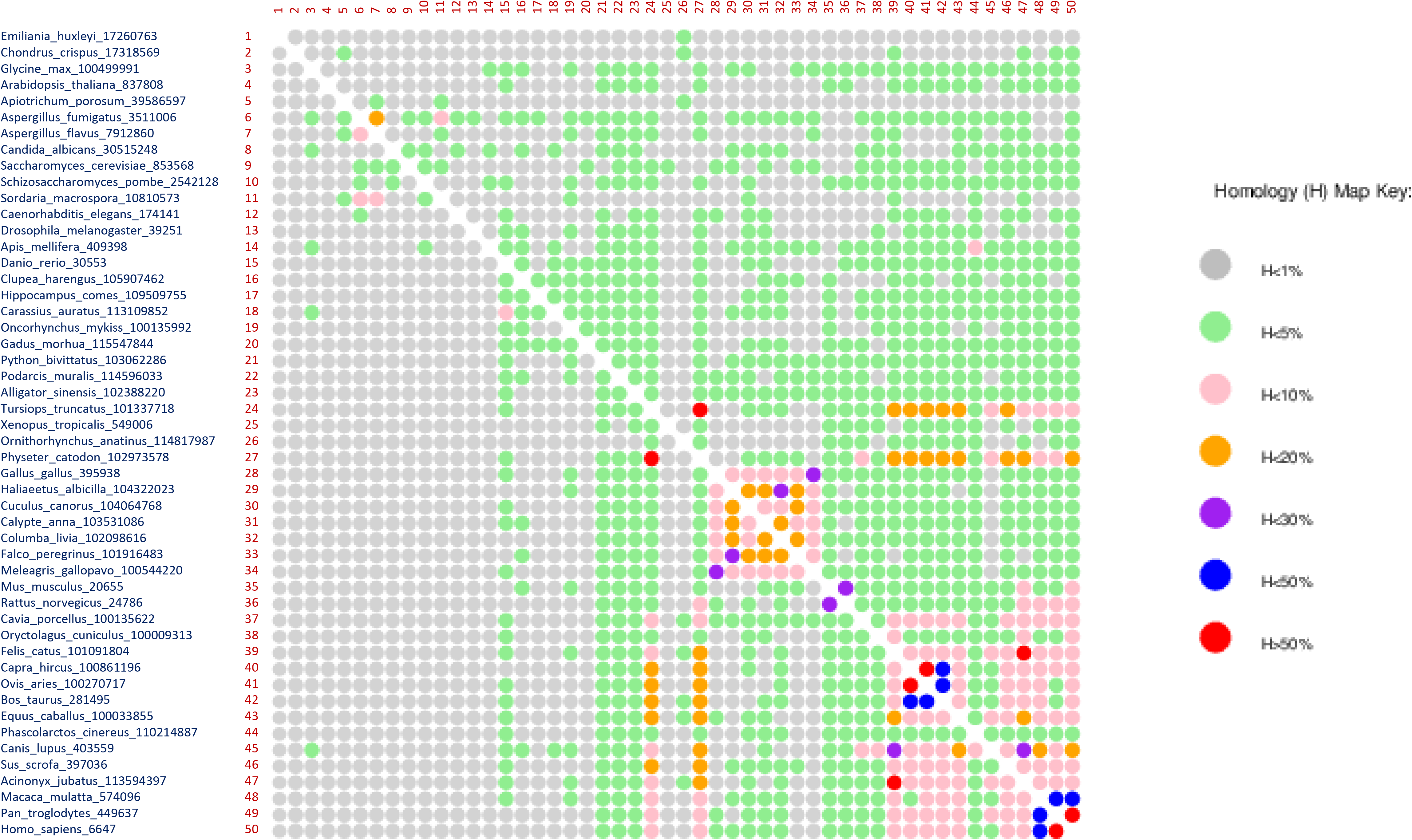
Homology Map of oligomer sequences across *SOD1* taxonomy. The map shows percentages of 10-mers in a *SOD1* gene that matched with its orthologs. E.g. there was >50% homology of *Homo sapiens SOD1* with *Pan troglodytes*. The map clearly shows that organisms lower in taxonomy contributed minimal (1-5%) k-mers to *SOD1* of higher organisms and that significant homology existed in narrow taxonomic groups only. This data is further detailed in Table S2.

### Effect of duplications and point mutations on LRC

In 48 of the 50 *SOD1* orthologs, the frequency of unique 10-mers was >90%. Hence, duplications were not in substantial frequency in *SOD1* in majority of organisms (Table S3). Only Two species, *Xenopus tropicalis* (549006) and *Ornithorhynchus anatinus* (114817987), had a frequency of repeated 10-mers ~ 20%. *X. tropicalis* had low fractal correlation (α=0.52) in spite of a significant gene length and multiple 10-mer duplications (Table S1 and S3). *O. anatinus* had high LRC with α=0.70, yet *D. melanogaster* (39251) had no duplications and still had an α=0.59 and *Rattus norvegicus* (24786) with unique 10-mer frequency of 99.3% had an α=0.68 (Table S1 and S3). These findings negate the view that segmental duplications are responsible for the LRC and fractal organization of genetic sequences. Moreover, 10 point mutations (A4V, D90A, E21G, G37R, H46R, I113T, I149T, L144F, L126Z and N65S) that cause ALS failed to individually alter the LRC characteristics of human *SOD1*. All 10 mutations tested together (which is a clinical impossibility), also did not affect the fractal measures (α=0.74, D=1.23).

### Reconstruction of *SOD1* sequences by randomization

If there were a 1000 species like *Drosophila melanogaster* with similar *SOD1* genes they would independently contribute only an average of 10 matching k-mers of length 10 nucleotides to Human *SOD1* (Table S4). In fact, natural *SOD1* orthologs provided comparatively greater number of matching 10-mers (Table S4). Theoretically, such randomly generated genes could *together* make 70% of Human *SOD1* sequence. As shown in Table S4 there is a linear relationship between gene length and number of random oligomers generated that will match Human *SOD1* sequence. In essence, a randomly generated sequence of over a million nucleotides will be able to provide all unique 10-mers in Human *SOD1*. Practically, however, this is not the case with majority of *SOD1* orthologs generated over millions of years of evolution providing <10% matching 10-mers (Table S3; Table S4). Random sequences with LRC were generated with CorGen for all *SOD1* orthologs. These random sequences could not replicate the natural *SOD1* orthologs and contributed minor frequency of 10-mers (Table S5) despite having α>0.5 (Table S1).

### Effect of fractal organization on Phylogenetics

Phylogenetic analysis showed that *SOD1* evolved in multiple clades (Figures 3, S2). DFA of 50 *SOD1* orthologous gene sequences showed that LRC were present in nearly all species (Table S1). Though there was general increase in LRC with taxonomic order as measured by α-values, the linear regression was poorly correlated (R^2^ = 0.49). However, when taxonomic order was rearranged according to α-values, R^2^ increased to 0.96 and 9 small taxonomic groups emerged that could be merged into 3 larger groups (Figure S3, S4). Phylogenetic analysis of these 3 groups showed *Emiliania huxleyi SOD1* (Gene ID: 17260763) which appeared to be initially closely related to *Xenopus tropicalis* (Gene ID: 549006) switched its phylogenetic position to be closer to Meleagris gallopavo (Gene ID: 100544220) and *Hippocampus comes* (Gene ID: 109509755) (Figure S4a). The group that remained stable was that of higher organisms (Figure S4c). The alteration of phylogenetic hierarchies according to DFA indicated that statistical topographies can be used to study gene evolution.

**Figure 3.**
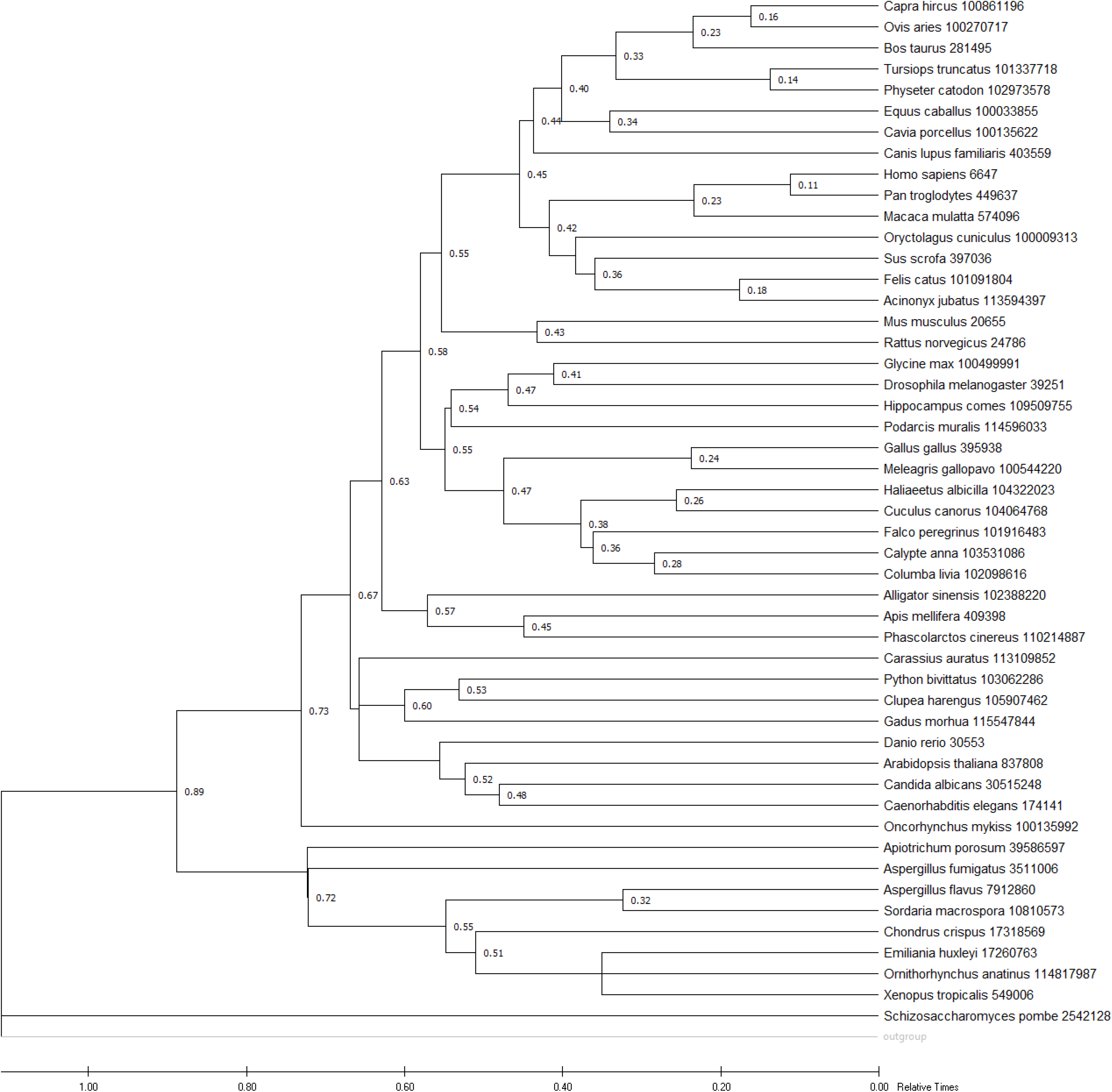
Time tree evolutionary analysis of *SOD1* orthologs. The timetree shown was generated using the RelTime method in MEGA X. Divergence times for all branching points in the topology were calculated using the Maximum Likelihood method and General Time Reversible model. The estimated log likelihood value of the topology shown is −348429.53. The tree is drawn to scale, with branch lengths measured in the relative number of substitutions per site. This analysis involved 50 nucleotide sequences. *Saccharomyces cerevisiae* (Gene ID: 853568) was used as outgroup.

## Discussion

This study showed that both short and long range statistical correlations exist in DNA sequences across taxonomic order. These correlations signify patterns in genomic organization that can be visualized graphically (Figure 1). The results here, point to the lack of emergence of such natural patterns from segmental duplications and point mutations. Though these fractal patterns track phylogenetic organization based on morphology, the minimal oligomer contribution across taxonomy indicated that *SOD1* gene did not evolve by a patchwork of sequences from simpler organisms.

Segmental duplication but not simple random sequence generation, has been postulated to replicate the frequency distribution of oligomers in ‘genome-as-text’ studies (Searls, 2002; Hsieh et al., 2003; Messer et al., 2005). In spite of statistical similarity between simulated and real genomes the sequence content was found to be widely discordant and failed to align (Hsieh et al., 2003). The natural sequence of *SOD1* orthologs could be theoretically reconstructed by an infinite number of iterations generating k-mers and patching them *together* in an infinite number of permutations – a process requiring infinite energy. In reality, nature conserves energy and as shown here, random processes were neither able to generate statistical correlations nor contribute substantially to the natural sequence of any of the *SOD1* orthologs (Table S3; S4; S5).

Previously, a limited study investigating the role of LRC in gene evolution showed that DFA α increased with taxonomic order of the *Myosin H* gene (Buldyrev et al., 1993). The study recommended that ideally precisely similar genes from different species should be analyzed, as has been comprehensively done in the present study for *SOD1*. Moreover, it is shown here that phylogenetics were altered due to fractal characteristics of gene sequences (Figures S3, S4). Thus, genetic LRC analysis may be utilized as a complementary method of studying gene evolution.

Pattern self-similarity observed here (Figure 1), has previously been attributed to sequence duplications (Albrecht-Buehler, 2012) which is refuted here based on the fact that *SOD1* orthologs had overwhelmingly (>90%) unique k-mer sequences. Point mutations have been thought to influence LRC (Li and Kaneko, 1992; Albrecht-Buehler, 2012). When 32 mutations in *SOD1* were evaluated in ALS patients, it was shown that they occur on unique haplotypes (Saeed et al., 2009), precluding the possibility of their natural coexistence. Yet, 10 *SOD1* point mutations *together*, also did not affect LRC and fractal measures of Human *SOD1*, indicating that point mutations likely play a minor role in LRC development.

What then could cause these correlations to exist? As shown in Table S3, *SOD1* orthologs had predominantly unique oligomer sequences. *Drosophila melanogaster SOD1* had no duplications, yet it showed substantial fractal organization (α = 0.59; D=1.31), while Human *SOD1* had 93.4% unique oligomers and had an α = 0.74. Not only were these oligomers unique within organisms, they were unique across taxonomic order (Table S2, Figure 2). These findings make obvious the fact that *SOD1* did not predominantly evolve as a patchwork of repetitive sequences. Instead, it is postulated here that specific underlying processes, such as nucleotide ligation, polymerization, elongation and restriction, led to a regulated weaving of sequences that shaped LRC. These and similar processes at the basic sequence level are likely to lead to sequence generation that has a specific order, generating LRC and are functionally biased. This kind of deterministic evolution would be time and energy efficient, allowing rapid and repetitive development of orthologous genes.

This study proposes that *SOD1* likely evolved independently in multiple clades over time, suggestive of convergent evolution (Orgogozo, 2015; Gompel and Prud’homme, 2009). Decades’ long experimental evolution of *E. coli* has shown that independent lineages coexisted for thousands of generations and led to evolution of the same phenotypic trait from only a few mutations (Blount et al., 2012). These and other evolution experiments have found repeatability at various levels of the genotype-phenotype pathway (de Visser and Krug, 2014). It has been postulated that five parameters viz pleiotropy, epistasis, plasticity, strength of selection, and population structure shaped evolution in a repeatable fashion that can be, at least partially, predicted (Stern and Orgogozo, 2009; Orgogozo, 2015). Moreover, the expanding assembly of instances of phenotypic convergence (Map of Life, 2020) indicates that evolution repeats itself at all stages ranging from molecules to ecosystems. The possibility of repeated evolution is exciting because of its inherent potential to unravel the deeper structure of evolutionary processes as well as predicting disease causation at the genetic level more robust.

## Methods

### Fractal measures

Fractal analysis was conducted on 50 orthologous *SOD1* sequences to determine the presence of long range order and self-similarity. The nucleotide sequence was recoded as numbers (A=1, T=2, G=3, C=4) for this analysis. The resulting sequence was subjected to detrended fractal analysis (DFA) as previously described (Peng et al., 1992, Saeed, 2005). The fractal exponent alpha was used as a quantitative measure. *SOD1* sequences from various species and randomly generated sequences were similarly assessed by DFA.

Relative dispersion analysis (RDA) employs Mandelbrot’s equation (ln L[l] = 1-D |n | + c) to quantitate the self-similarity and is based on the box-counting method (Glenny et al., 1991; Saeed, 2005). Plotting the variation in the signal against the window size on a log-log scale quantifies the fractal dimension (D) and is evidence of power law behavior (Peitgen et al. 2010).

The RDA principle was used to generate a novel graphic representation of the genetic sequences to evaluate scale invariant pattern formation i.e. whether the basic architecture of short sequences in a given *SOD1* gene is similar to that of its complete sequence. Nucleotides A, and T were plotted in the vertical plane and G and C in the horizontal plane. Shorter sequences with 2^n^ nucleotides (8 ≤ n ≤ 13) were drawn in proportional window sizes to visually evaluate self-similarity, as in zoomed-in photographs. The programming for all methods described here was carried out in Python 2.7.9 and incorporated in the software, *GeneFractals*.

### k-mer restriction

Frequency of a k-mer is the count of its occurrence in a sliding window of width k, as the window moves across the gene once (Hsieh et al., 2003). Sliding window analysis was performed for reference *SOD1* gene sequences with a 10-nucleotide window size (k=10). This allowed the *SOD1* sequence to be chopped up into N overlapping 10-mers depending on the length. The frequency of the 10-mers was subsequently compared with *SOD1* from other species or randomly generated sequences to quantitate similarity. The frequency matching across species was visualized in a graphic table.

### Random sequence generation

Frequency analysis of randomly generated short k-mers was used here as in other molecular evolution studies (Hsieh et al., 2003). However, the k-mer model does not have long-range correlations in the sequence composition. To compensate for this deficiency the dynamic model incorporated in CorGen was employed (Messer and Arndt, 2006). CorGen can generate random sequences with the same correlation and composition parameters as the actual input sequence (e.g. mouse *SOD1*). Hence, both models were used to generate random sequences that were tested for base composition similarity with real *SOD1* gene sequences across a variety of species.

### Phylogenetic analysis

Complete genomic sequences of 50 *SOD1* orthologs were retrieved in FASTA format from NCBI through Gene search. The sequences were aligned with the MUSCLE algorithm incorporated in MEGA X software (Kumar et al., 2018). Construction of phylogenetic trees and estimation of evolutionary distances for all sequences was performed using the Maximum Composite Likelihood method in MEGA X (Tamura et al., 2012).

## Acknowledgement

I am grateful to Aneela Pasha for her insightful comments on the manuscript and valuable discussion and encouragement throughout this project.

Competing interests: None

## Data availability statement

All data relevant to the study are included in the article or uploaded as supplementary information. All datasets are in publically available repositories with their accession numbers in the manuscript.

## Web Resources

*GeneFractals* software will be made available on GitHub subsequently GeneFractals SOD1 Images: https://drive.google.com/drive/folders/1XPFG8FTtOZAet-saFOQM34Vo3WVfq_eT?usp=sharing SOD1 mutations causing ALS and their genomic positions were obtained from ALSoD database: https://alsod.ac.uk/output/gene.php#geneSummary

**Figure. S1. Varying self-similarity of *SOD1* orthologs** The GeneFractal diagrams for *SOD1* sequences of *Apis mellifera*, *Arabidopsis thaliana* and *Homo sapiens* are shown. They demonstrate varying visual self-similarity and fractal measures as indicated.

**Figure. S2. Phylogenetic analysis of *SOD1* orthologs** The evolutionary history is shown as an unrooted tree of 50 *SOD1* orthologous sequences. It was inferred by using the Maximum Likelihood method and General Time Reversible model in MEGA X. Initial tree(s) for the heuristic search were obtained using Neighbor-Join and BioNJ algorithms to a matrix of pairwise distances estimated using the Maximum Composite Likelihood (MCL) approach, and then selecting the topology with superior log likelihood value.

**Figure. S3. *SOD1* Orthologs graphed by DFA** The *SOD1* orthologs were ordered according to increasing DFA α-values. The graph shows the rearrangement of orthologs in 9 groups according to their varying gradients. These groups (A-I) were further grouped into 3 larger groups (Groups D, E, F were classified as Group 2) that were used for phylogenetic analysis in Figure S4.

**Figure. S4. *SOD1* Clades according to DFA based fractal organization** The evolutionary history of *SOD1* orthologs (n=15) was evaluated with MEGA X, using the same processes as in Figure S2. a) In this unrooted tree (Group 1), *Emiliania huxleyi* switched its position from the group of *Xenopus tropicalis* (Figure S2) to that of *Meleagris gallopavo*. This shows that fractal correlations can be used to study phylogenetic organization. b) There were 4 clades in Group 2 in line with careful observation of the gradients in Figure S3. c) Group 3 containing mammals remained unchanged from Figure S2.

**Table S1. Fractal measures of *SOD1* orthologs** Shows the species name (n=50), their *SOD1* NCBI gene ID, NCBI reference sequence ID, NCBI taxonomic order, DFA Groups according to Figures S3 and S4, percentages of AT and GC content and the DFA of CorGen generated random sequences with the natural *SOD1* ortholog as the input sequence.

**Table S2. Phylogenetic sequence reconstruction of *SOD1*** Shows the calculated frequency and number of matched k-mers (k=10) across 50 *SOD1* orthologs. The highlighted species_gene ID sequence was analyzed by sliding window analysis and used as a reference for matching the k-mers generated by similar analysis of all 50 orthologs (testing sequences). The percentage match refers to the number of k-mers in the reference sequence that matched the testing orthologous sequence.

**Table S3. Analysis of natural and random oligomers for *SOD1* orthologs** Sliding window k-mer (k=10) analysis of natural *SOD1* orthologs produced predominantly unique k-mers. E.g. *Emiliania huxleyi* (Gene ID: 17260763) with gene length of 658 bp, produced a total of 555 k-mers, of which 530 were unique (95%), 1 k-mer occurred 9 times and 2 k-mers occurred 8 times in the sequence. Randomly generating 555 k-mers (k=10) over 10 cycles, led to a total of 43 k-mers that matched the natural k-mers of *Emiliania huxleyi*. Each cycle generated a mean of ~4 matching k-mers (range 2-6).

**Table S4. Reconstruction of Human *SOD1* sequence by simple random oligomer generation** *Homo sapiens SOD1* (Gene ID: 6647) was subjected to sliding window k-mer (k=10) analysis. *SOD1* orthologs from 49 other species were similarly analyzed and their k-mers matched with the k-mers of Human *SOD1*. Random k-mers were generated over 10x cycles and the mean number per cycle was used for calculating the odds ratio. All natural sequences contributed higher frequency of matching k-mers to Human *SOD1* than randomly generated sequences. Three species were exceptions (*Sordaria macrospora*, *Emiliania huxleyi* and *Apiotrichum porosum*), indicating that their contribution could be due to chance. The experiment was repeated for limited number of species (due to computing time required) for 100x and 1000x cycles however, the mean number of matching k-mers remained the same as for 10x cycles. This process mimicked increasing the gene length and number of random matching k-mers could be calculated using the equation: Number of Random 10-mer = 0.0062 × Gene length (nucleotides); R^2^ = 0.99.

**Table S5. Matching complex random oligomer generated using CorGen with natural *SOD1* orthologs** CorGen generated random gene sequences (prefixed with ‘r’) matched with random and natural *SOD1* orthologs by k-mers analysis as previously described. Minimal k-mer matching was produced for natural *SOD1* sequences.

